# The hidden predictors of human haematopoietic clonal fate

**DOI:** 10.1101/2025.07.28.667115

**Authors:** Sara Tomei, Stephen Zhang, Michael Lin, Cindy Audiger, Tom Weber, Shalin H. Naik

## Abstract

Human haematopoietic stem and progenitor cells (HSPCs) exhibit heterogeneous lineage output, but the molecular programs underlying clonal fate remain poorly defined. To address this, we developed a human haematopoietic organoid supporting differentiation into 15 lineages and used it to track barcoded HSPC clones over time. By integrating single-cell transcriptomes, surface phenotypes, and clonal fate, we applied machine learning to identify *clonal fate modules* – gene and marker signatures predictive of lineage commitment. This approach uncovered hidden transcriptional and surface correlates of multipotency, including CD200, which marked a subset of HSCs with broad output capacity, which we leveraged to increase manufactured type 1 dendritic cell purity for immunotherapy applications. Our study provides a framework for decoding clonal fate decisions in human HSPCs and identifies molecular features that distinguish truly multipotent clones, advancing strategies for stem cell purification and therapeutic engineering.

## Introduction

Haematopoiesis is the highly regulated and complex hierarchical system of red and white blood cell production and maintenance. At the apex reside hematopoietic stem cells (HSCs), which possess the unique capacity for both self-renewal and differentiation into all blood cell lineages. These HSCs progressively give rise to committed progenitor populations, which ultimately differentiate into mature blood lineages. Advances in single-cell ‘omics including RNA sequencing (scRNA-seq), CITE-seq and ATAC-seq have enabled a deep characterization of these haematopoietic stem and progenitor (HSPC) populations providing a detailed map of the stages and trajectories of hematopoietic development ^1-7^.

However, whether such ‘maps’ best reflect haematopoiesis at a clonal level is challenged by the numerous observations that, even within transcriptomically and phenotypically similar populations, HSPCs can be highly heterogeneous in their clonal fate^8-16^. Furthermore, the molecular determinants underlying many of these diverse fate choices remain unknown. This raises an open question: can one predict clonal behaviour using common approaches such as single cell ‘omics with unbiased clustering? Or are the predictors of clonal fate “hidden” such that each single cell’s fate must also be explicitly known to extract the correlated molecular features?

If the latter, this poses a challenge considering the same single cell cannot be tested for both its molecular profile (single cell ‘omics is destructive) and its fate (the founder cell is lost through division and differentiation). The advent of clonal multi-omics – where single cells are induced to divide and the clonal siblings tested in separate assays – allows a practical solution to this challenge. SIS-seq^17,18^ and similar methods^16,19-23^ allow these clonal siblings to be tested in parallel SISter assays of fate and scRNA-seq and have been successful in unearthing novel regulators of clonal behaviour in haematopoiesis, reprogramming, development and cancer.

In haematopoiesis, clonal multi-omics has largely been applied in murine studies in vivo^16^ or in vitro^18,22,24^, but not yet for human haematopoiesis. Examining clonal fate in human haematopoiesis could conceivably utilise synthetic or natural barcodes, and be tested i) in vitro, ii) in humanised mice, iii) in humans after stem cell transplantation, iii) or in humans directly. While the last three provide the most in vivo relevant models, they can be logistically difficult, cost prohibitive, and limiting in terms of experimental possibilities. Instead, in vitro models allow a more tractable system, including the study of clonal dynamics with high recovery of progeny compared to serial bleeding in transplanted mice or humans, for example. Unfortunately, most models of human haematopoiesis generate only a few of the multitude of haematopoietic cell types ^1,15,23,24^. A putative human haematopoietic organoid using CD34^+^ progenitors as starting material that is genuinely multi-lineage, and reflects cell diversity across multiple organs, would facilitate a clonal multiomics approach of human haematopoiesis.

Here we present an in vitro organoid of human haematopoiesis that enables the production of up to 15 different lineages in a single vessel. By coupling this with lentiviral barcoding, clonal multi-omics and machine learning analysis, we identify the hidden clonal features of human hematopoietic differentiation and validate CD200 as a novel multilineage HSC marker, from which most dendritic cells (DCs) are produced.

## Results

### A multilineage human haematopoietic organoid

Determining which cell types an HSPC clone can produce (i.e. its potency/potential) requires an assay where all fates are possible. This is problematic in differentiation assays that only favour one or a few lineages. This also applies after adoptive transfer in vivo, where a clone’s fate could be determined by the niche it lands in. To this end, we developed a human haematopoietic organoid system we term Haematopoiesis using cyTokines and Optimised Niches In Culture (HaemaTONIC). It involved the co-culture of human CD34^+^ progenitors with murine stromal cell lines supplemented with a cocktail of cytokines (Figure 1a).

**Figure 1.**
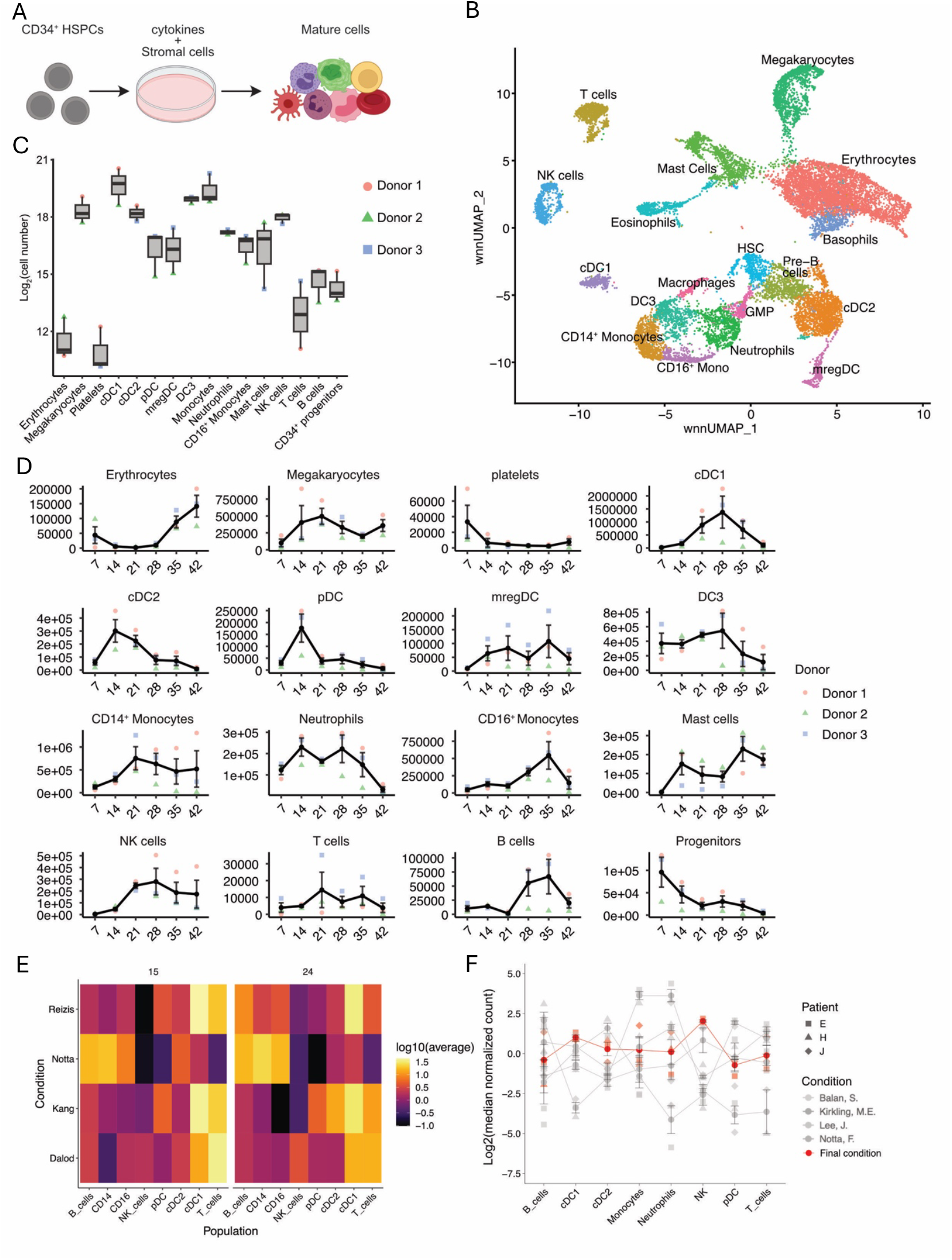
HaemaTONIC permits multilineage human haematopoiesis in a single well. a) HaemaTONIC culture system. b) WNN UMAP using transcriptome and surface marker expression (TOTAL-seq) of cord blood-derived cells after culture in HaemaTONIC (data from 6 different donors with pooled output at day 7 and day 21). c) Enumeration of cell types developed in HaemaTONIC culture at day 21 with spectral cytometry panel. d) Number of cells from each lineage produced throughout the culture. **e)** Heatmap showing number of cells produced in other protocols (average of 3 donors) at day 15 and day 24 **f)** Number of cells produced in each protocol normalised to average of cells produced in all protocols (HaemaTONIC in red)

This system was able to give rise to at least 15 different cell types as measured by scRNA-seq + TOTAL-seq (Figure 1b). This included the development of erythrocytes, megakaryocytes, monocytes, neutrophils, eosinophils, basophils, mast cells, plasmacytoid DC, cDC1, cDC2, DC3, mregDC, NK cells (CD56 dim and bright), B cells and T-like cells (T cells cannot develop without thymus-like selection). These cells were highly transcriptionally similar to their corresponding lineage in a reference human bone marrow dataset (Supp Fig 1a-b). Our TOTAL-seq data set was used to inform the design of a multiparameter spectral cytometry panel which we used to validate HaemaTONIC output with flow cytometry (Figure 1c-d, Supp Fig 1c). We tracked the culture across 6 weeks and assessed that HaemaTONIC was robustly able to support multilineage haematopoiesis across multiple donors (Figure 1c). We noted that different populations were produced with different kinetics, with myeloid cell production peaking around day 14-21 and lymphoid and mast cell production peaking at around day 35.

Through comparative analysis, we noted that existing protocols^11,15,25,26^ produced differing numbers of each cell type and none was able to produce all the human cell types that HaemaTONIC could (Figure 1d-e). Notably, the Lee protocol favoured production of DCs (cDC1, cDC2 and pDC), the Lee and Balan protocols yielded the most pre-T cells, and the Notta protocol was the most efficient in producing neutrophils, monocytes and B cells. The Notta protocol also generated cDC2s, comparable with Lee, but not cDC1s. Notably, none of the protocols were able to generate substantial amounts of NK cells, indicating that none of these conditions were suitable for this cell type. HaemaTONIC therefore demonstrated superior performance in both lineage diversity, ‘balance’ of cell production, and was composed of cell types found across multiple organs: PBMCs (enriched for lymphoid and myeloid cells), BM (enriched for HSPCs, megakaryocytes, erythroid progenitors), and spleen (enriched for the different DC subtypes). For these reasons, we pursued HaemaTONIC as a system to identify the features of human clonal HSPC fate.

### Clonal fate is conserved in human HSPCs

Beginning with CD34^+^ cord blood progenitors from six donors, we uniquely labelled each with one of three different SPLINTR barcode libraries^20^, each coupled to a distinct fluorescent protein: GFP (donors 1 and 2), BFP (donors 3 and 4), or mCherry (donors 5 and 6) (Figure 2a). These libraries were distinguishable either by flow cytometry or through bioinformatic barcode analysis, enabling the multiplexing of at least three donors together.

**Figure 2.**
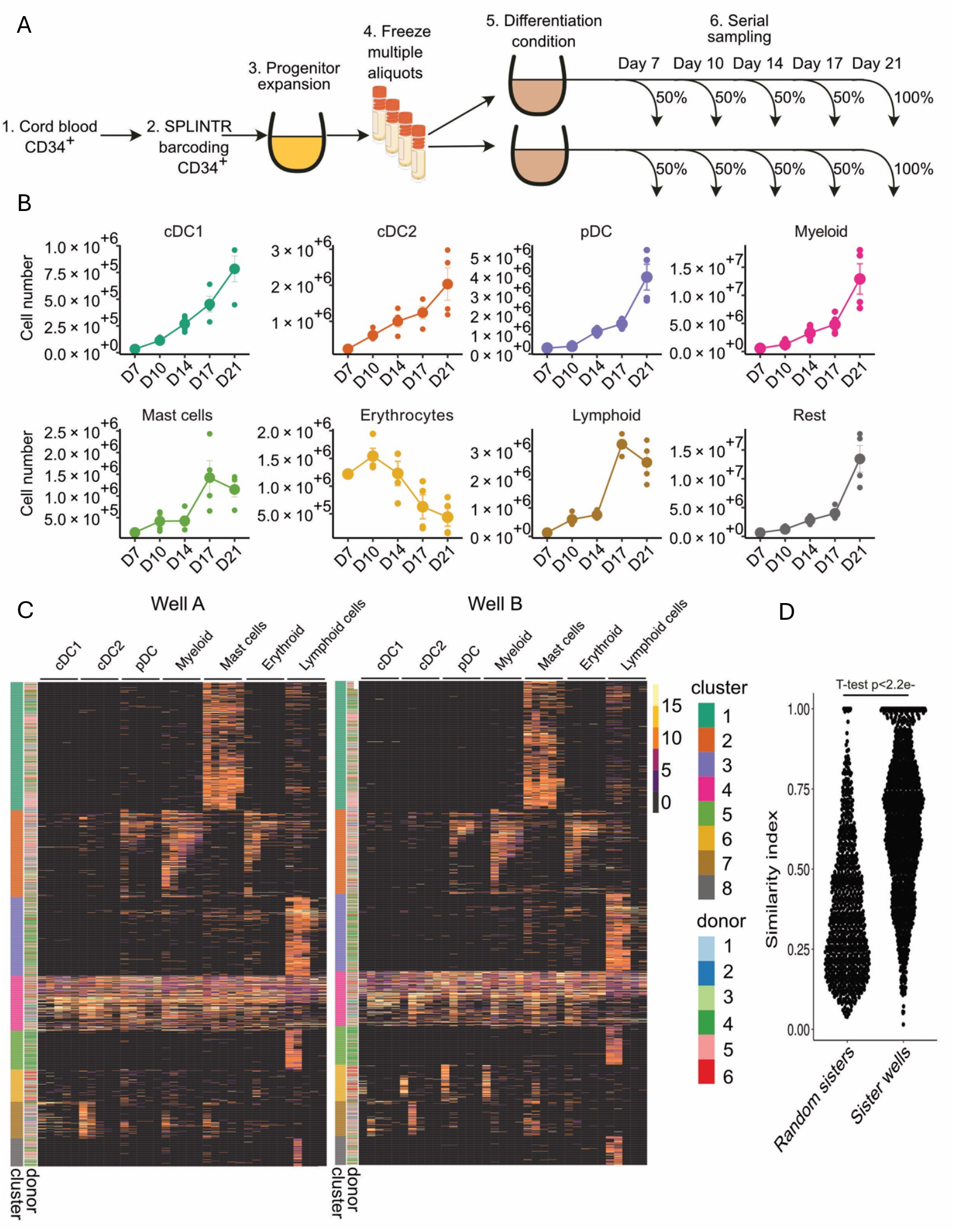
Intrinsic programming governs lineage output in HSPCs. **a)** SIS-seq experimental layout. Barcoded cells were expanded and then frozen in multiple aliquots each containing a few copies of each barcode. The aliquots were used to test conservation of fate when cells were cultured in separate wells. **b)** Cell number for populations sorted for barcode analysis. **c)** Heatmap showing clonal fate in both sister wells. Cell population annotated on top, each column is a different timepoint (from left to right: day 7, day 10, day 14, day 17 and day 21). **d)** Similarity index between sister wells.

Following labelling, CD34-enriched progenitors from each donor underwent expansion in StemMACS media (Figure 2a). When cultures reached between 4-5 × 10^6^ cells in total, or day 11 of expansion, whichever came first, cells were frozen down in aliquots of 5 × 10^5^ cells each (Figure 2a). Two aliquots were thawed and progenitor cells cultured on HaemaTONIC in 2 different wells, and a third aliquot was defrosted for scRNA-seq + TOTAL-seq (Figure 2a, data in Figure 3).

**Figure 3.**
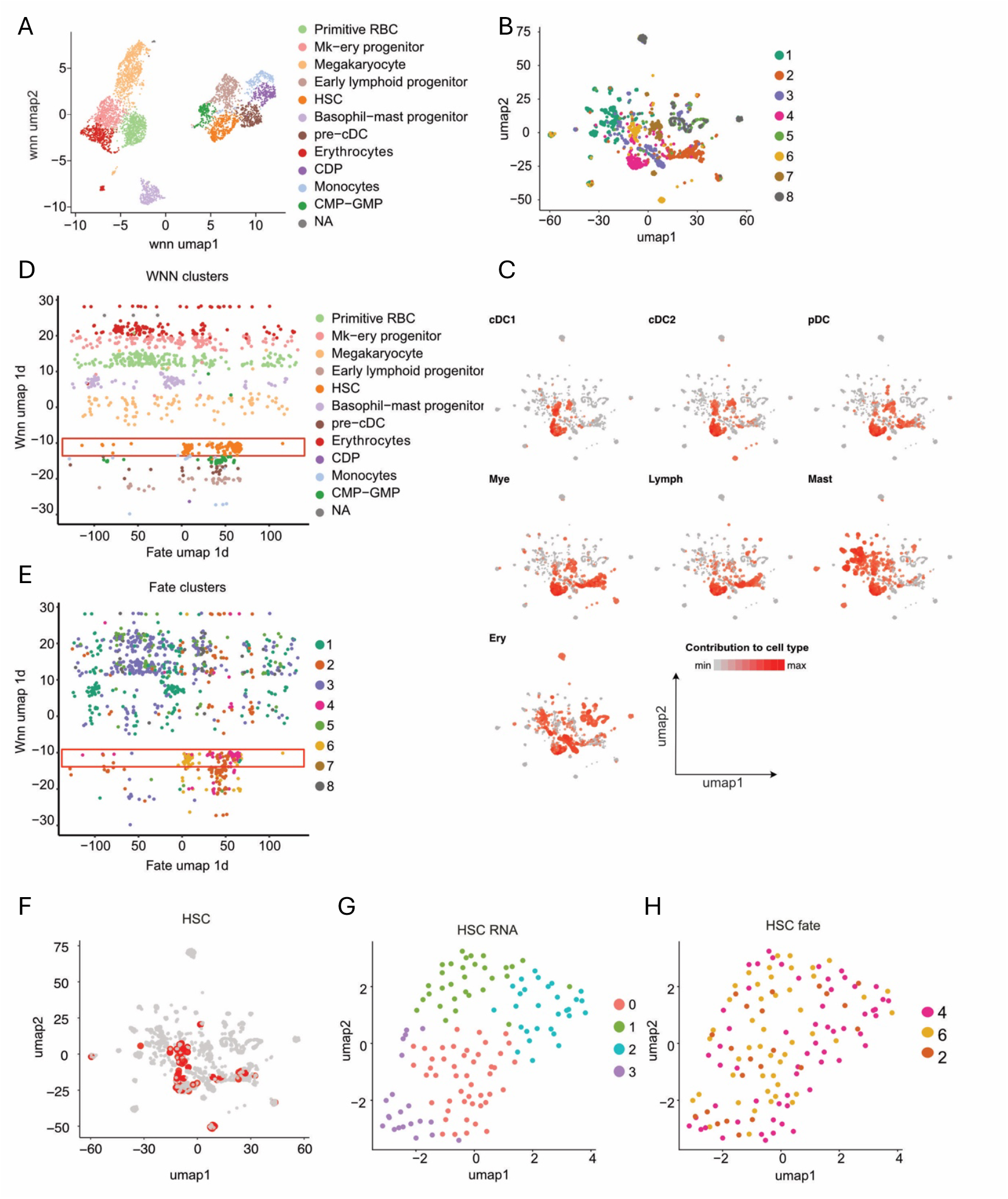
Unbiased single cell ‘omics does not predict clonal fate. **a)** WNN UMAP of CD34^+^ HSPC after the expansion phase (10 days). **b)** Fate UMAP on bulk barcode data obtained from mature populations sorted from HaemaTONIC (Fig. 2) **c)** UMAP from b) overlaid with lineage fate from any time point. Colour scale is the log_2_ of biomass of the indicated population at the timepoint analysed (one dot per timepoint per barcode) **d)** One-SENSE visualisation of cells: WNN 1D UMAP from a) on y-axis and the 1D UMAP of clonal fate from b) on x-axis. Cells are coloured by d) WNN clusters or **e)** by clonal fate cluster. **f)** Clones highlighted on the fate UMAP from b) whose siblings were in the HSC cluster from a). **g)** Sub-clustering of the HSC transcriptional cluster based on transcriptome and surface marker expression profile alone, without clonal fate. Clones are coloured based on their **g)** Seurat cluster **h)** or fate cluster.

However, an important pre-requisite for SIS-seq in every new setting is to first test whether clones are similar in their fate (conserved) in replicate sister assays. When a feature is clonally conserved, it reflects a heritable program that could be measured in the clonal siblings. To test this, half the contents of each replicate well was harvested every 3-4 days starting from day 7 to test the barcode repertoire in the resulting progeny (Figure 2a). For this analysis we tracked output from Day 7 to day 21 as it captured most lineages (Figure 1d). This would allow the testing for conserved fate as well as recording HSPC clonal dynamics.

We developed a step-wise enrichment, cytometric staining panel, and 4-way sorting strategy (Supp Figure 2a) to recover 8 populations for barcode analysis at these time points: cDC1s (CD1c^+^Clec9A^+^), cDC2s (CD1c^+^Clec9A^−^), total Myeloid cells (CD1c^−^ (CD14^+^ and/or CD16^+^)), pDCs (CD1c^−^CD14^−^CD16^−^CD123^+^), Erythrocytes (CD45^−^GlyA^+^), Mast cells (CD45^+^cKit^high^), Lymphocytes (CD7^+^ or CD19^+^ cells), and ‘Rest’ (CD45^+^ cells that did not stain for any of the markers analysed) (Supp Figure 2b). In this way, while some populations were pooled for practicality, no cells were excluded from analysis. We opted for population-based barcode analysis as it is more cost effective, and more sensitive in detecting small clones amongst large numbers of cells, compared to assessing fate through single cell analysis. We acknowledge this provides less resolution to measure differential contribution to cell subsets and states. Each population was then lysed and DNA was extracted for barcode PCR and sequencing followed by computational analysis to determine the clonal contribution of different donor HSPCs to the different cell types.

The culture system successfully generated all lineages, albeit with different kinetics (Figure 2b). Erythrocytes developed early, peaking at day 10 while mast cells and lymphoid production remained quite low and peaked later at day 17. In contrast, DCs (cDC1, cDC2 and pDC) and Myeloid cells exhibited continuous increases until the end of the culture period. HSPCs cultured in different wells from the same donor produced the same number of progeny cells at each timepoint (Figure 2b).

We then appraised the clonal contribution of each progenitor in HaemaTONIC. Over 3000 barcodes were recovered spanning all time points across the 6 donors (Supp Figure 2c). Of these, 44% were found in at least one cell type in at least one time point, in both replicate sister wells (Supp Figure 2d). These clones accounted for over 90% of all cells produced (biomass) in each well (Supp Figure 2e). This suggested the remaining 56% of barcodes in only one well largely represented small clones that were either below the detection threshold in the sister well, or absent due to their small size during clone splitting. Filtering the dataset for common barcodes between wells allowed us to retain clones that contributed to most of the biomass.

Clones expanded and contracted with different dynamics. The same clone cultured in separate wells followed very similar dynamics, indicating that clonal waves were conserved between sister wells (Supp Figure 2f). This feature was observed across donors. Further, we assessed the output of each clone at every timepoint (Figure 2c). Remarkably, even the type of cells that each clone produced at each timepoint was mostly conserved between wells (Figure 2c) suggesting that human HSPCs are imprinted not only for what lineages they will produce, but also the kinetics and quantity of that production, similar to mouse HSPCs^24^.

We clustered clones (a 1D UMAP based ordering and clustering of clonal fate heterogeneity) based on their production dynamics and identified eight broad clusters (Figure 2c). Cluster 1 was biased towards mast cell production across multiple timepoints. Clusters 3, 5 and 8 all produced erythroid cells with differing kinetics. Cluster 2 was myeloid biased with production of lymphocytes and cDC2 at earlier timepoints. Cluster 6 and 7 comprised barcodes that had an early wave of cDC1, cDC2, pDC, myeloid and mast cell production. A substantial multi-/oligo-potent cluster 4 was also identified, producing all/many lineages at nearly all timepoints. These fates were consistent with the notion that CD34^+^ HSPCs are heterogeneous at the time of isolation and barcoding.

In addition to visualising conserved clonal fate (Figure 2c) we could also measure it statistically (Figure 2d). Remarkably, a separate frozen aliquot of barcoded HSPCs from the same donor cultured several months later in HaemaTONIC retained similar clonal fate, demonstrating the robustness of both HSPC lineage priming and our culture system (data not shown). Importantly, the pre-requisites for SIS-seq were now met: lineage bias was demonstrably heritable, allowing us to leverage the siblings of these progenitors for determining the transcriptional and surface marker predictors of fate.

### Single cell ‘omics alone cannot predict fate

To identify these predictors, a separate aliquot from each of 4 donors was defrosted, stained with TOTAL-Seq A (a commercially available panel of 163 oligo-conjugated Ab), sorted for fluorescent protein-expressing (barcode^+^) cells, and captured using 3’ scRNA-seq (Figure 3a, Supp Figures 3 and 4 – heatmaps of Tc and Prots). The resulting data analysed using WNN separated in two main groups and sub-clusters using WNN (Figure 3a); the left contained erythrocyte (*Hbb, Hbg1, Hbg2*) and megakaryocyte progenitors (*Itga2b*), the right contained myeloid (*Mpo*), lymphoid (*Il7r* and *Ltb*), and DC progenitors (*Irf8*), primitive stem and progenitors (*Spink2, Msi2, Avp*), and a basophil-mast cell cluster (*Hpgd* and *Prg2*) (Figure 3a).

**Figure 4.**
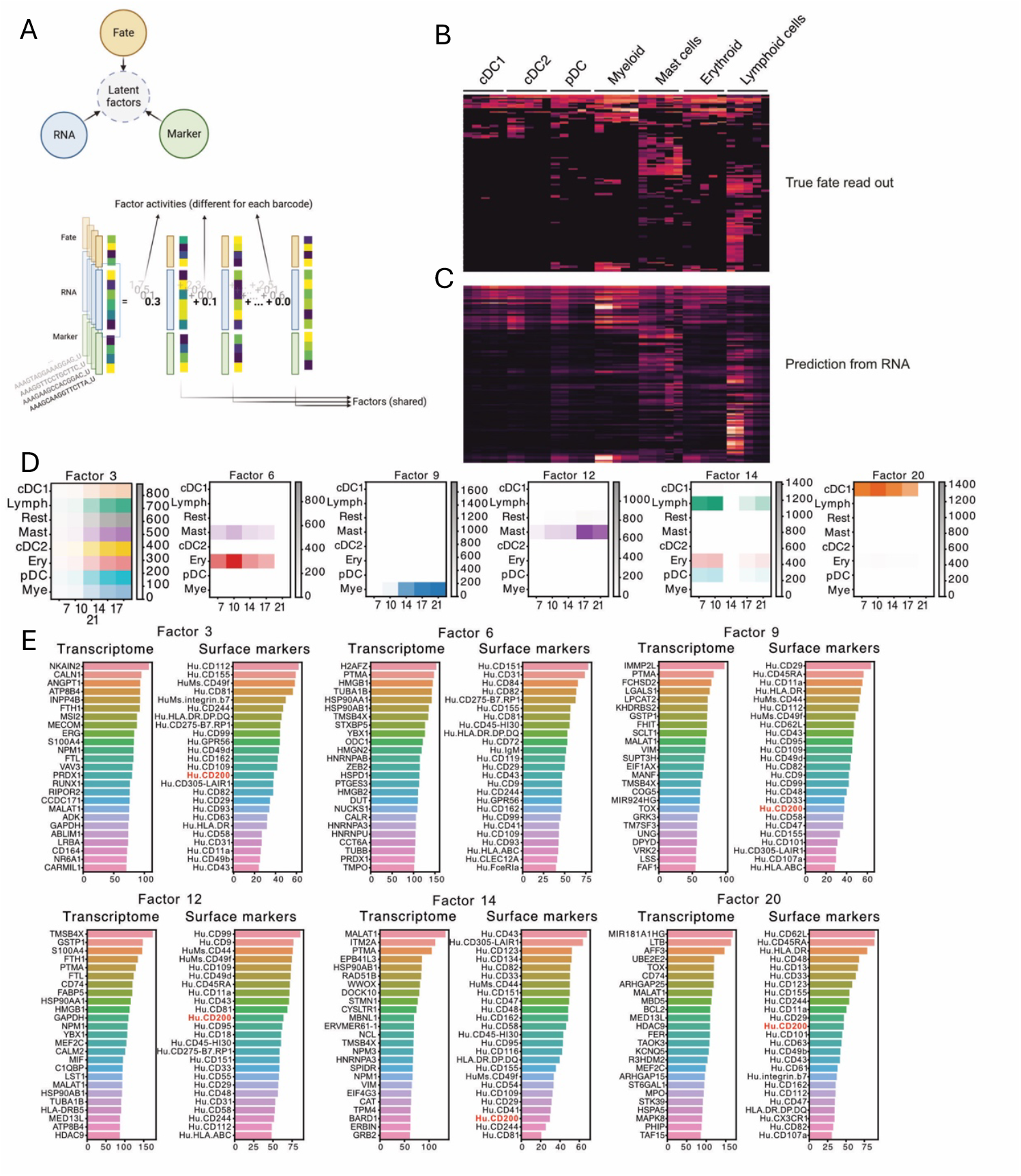
Factor analysis identify genes and markers that predict fate. **a)** Factor analysis layout. **b)** Barcode heatmap showing true fate outcome in HaemaTONIC and **c)** predicted fate outcome using only the RNA information. Each row is a population sorted at a specific timepoint. Columns are grouped by population (annotated on the top) and timepoint are ordered left to right (Day 7, 10, 14, 17, 21). **d)** Heatmap summarising the fate of each factor or CFM (clonal fate module). **e)** Genes and surface markers associated with each factor/CFM.

Having clustered the same cells according to both A) single cell ‘omics using WNN (Figure 3a) and B) clonal fate (Figure 3b, UMAP derived from Figure 2c), we could answer our original question: is ‘unbiased’ single cell molecular analysis alone (A) able to predict clonal fate (B)? This was first visualised using One-SENSE^27^, which allows orthogonal comparison of one measurement to another for the same single cell. For the 1,064 single cells that also harboured a barcode in the bulk barcoding time series data, One-SENSE visualisation of WNN data (y-axis) to clonal fate (x-axis) (Figure 3d-e), revealed numerous examples of poor molecular:fate concordance, both visually (Figure 3d-e). This was also visualised by superimposing clones from each WNN cluster on the fate UMAP. We did this for WNN-annotated “HSCs” (Figure 3f), which spanned different fates (clusters 2, 4 and 6), and all other progenitor populations (Supp Figure 5). To rule out whether this was due to poor clustering granularity, we extracted then clustered HSCs into 4 different clusters (Figure 3g). However, this did not better separate fate clusters of “HSCs” (Figure 3h).

**Figure 5.**
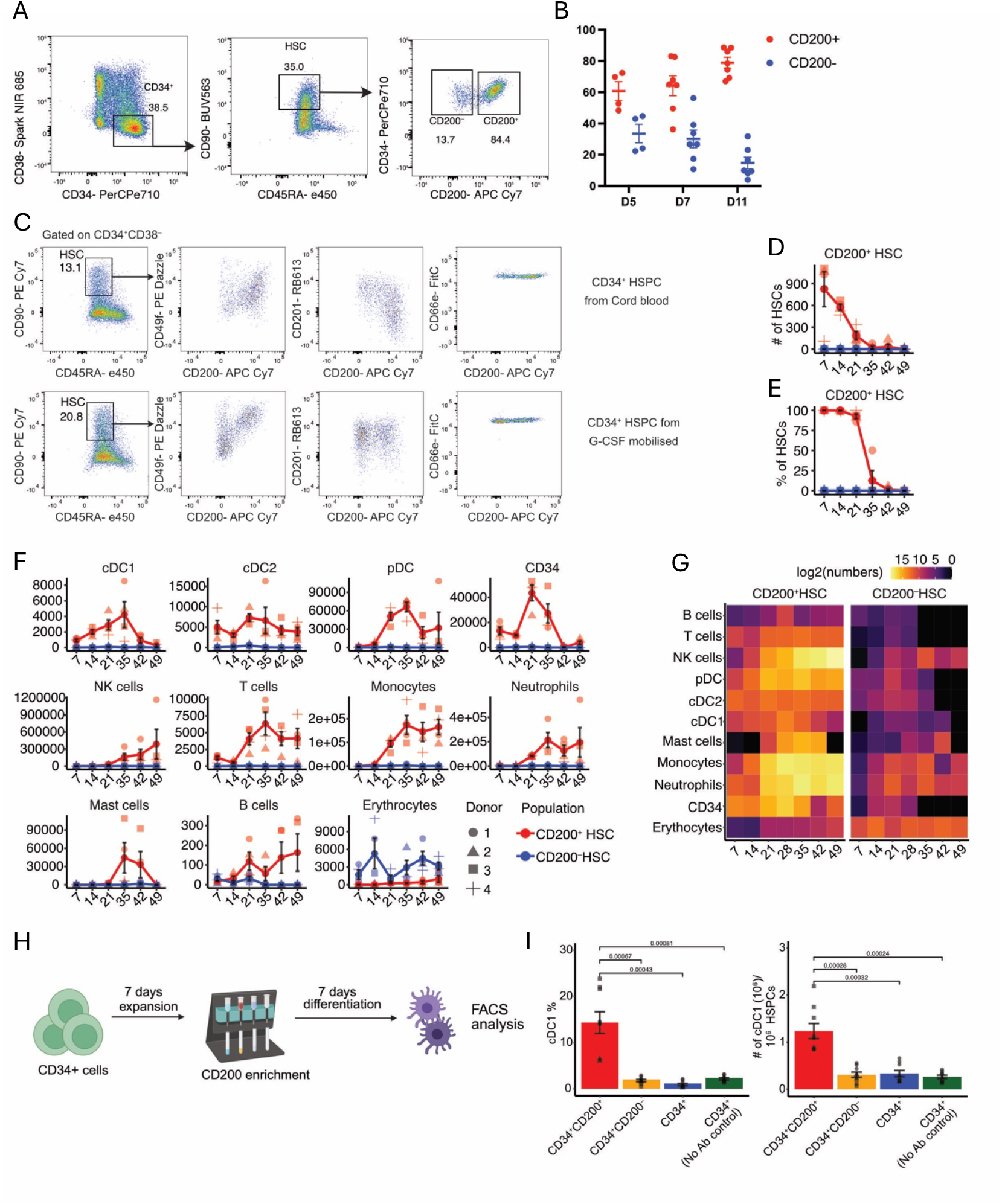
CD200^+^ and CD200^−^. HSCs are two functionally distinct populations **a)** Phenotypic characterization of expanded cord blood samples at Day 11. **b)** Percentage of CD200^+^ and CD200^−^ HSCs during the expansion phase in 4 cord blood and 3 G-CSF samples. **c)** Comparison of CD200 with other previously identified HSCs markers at day 5 of expansion. Top row shows 4 concatenated cord blood donors while the bottom row shows 3 concatenated G-CSF donors. **d)** CD200^+^ HSC numbers in HaemaTONIC. **e)** Percentage of HSC that are CD200^+^ in HaemaTONIC. **f-g)** Numbers of mature cells produced by CD200^+^HSC and CD200^−^ HSC in HaemaTONIC by 4 different cord blood donors. **g)** summary heatmap of f). **h)** Experimental layout of CD200 enrichment for GMP culture. **i)** cDC1 percentage and numbers produced in GMP differentiation condition

These analyses demonstrate that the HSPC marker genes and proteins that drive clustering are not necessarily the same as those that predict what cell types a stem or progenitor cell will make i.e. a functional definition. It was possible that either a feature that we did not measure controls fate (e.g. epigenetic modifications, metabolome, cell size or other) and/or only a subset of “hidden” gene and protein expression correlated with fate. While the former is conceivable and likely, we focused on the latter.

### Factor analysis identifies hidden clonal fate modules

We postulated that finding the molecular features of HSPCs predictive of fate first required incorporation of clonal fate into an analysis. To achieve this we opted for factor analysis: a machine learning algorithm used previously to integrate single cell and multi-omics data^28-30^. We reasoned it would be suitable for learning clonal multi-omics data and developed it to simultaneously incorporate the clonal dynamics of fate, transcriptome and surface marker expression of each clone to learn a series of latent factors that explain the variability (Figure 4a).

Prior to learning the factors, we applied two pre-processing steps. First, for clones that were represented by multiple cells with the same barcode (i.e. developed from the same ancestral cell), we averaged their transcriptome and surface marker expression. This pre-processing step ensured that each clone had a single unified value for each gene and surface marker associated with its clonal fate. We acknowledge biologically relevant information may have been lost through averaging. Second, the fate component included a data matrix that included each barcoded clone (rows), and its temporal contribution to each cell type (one column per cell type per time point).

A first analysis revealed 15 different factors, hereby referred to as clonal fate modules (CFMs), that sufficiently explained the dataset variability (Supp Figure 6). Each of these CFMs contained the associated i) clonal dynamics of fate, ii) genes expressed, and iii) surface markers (Figure 4a and Supp Figure 6). The components of each clone in the dataset (transcriptome, fate and surface marker expression) can then be scored against each factor. As a result, each clone harboured different levels of activity for each factor. It was re-assuring that each CFM identified known developmental genes, such as *MPO* and *SERPINB1* for myeloid development (CFM 1), *GATA2* and *KIT* for Mast cell development (CFM 2), *HBD* for erythroid cells (factor 9).

To test the robustness of the clonal fate modules, we trained the model on 80% of the clones and then tested whether the resulting gene expression values alone of each CFM, which were previously hidden, could now predict fate of the remaining 20% (Figure 4b-c). This approach successfully predicted clonal fates solely from CFM gene expression values (Figure 4b-c). The same was not true for the predictive value of the limited number, and surface expression bias, of proteins assessed using the TOTAL-seq panel (Supplementary Figure 6). Nevertheless, the ability to predict clonal fate from a selection, but not all, gene expression values from the CFMs suggests that *bona fide* predictors of fate do exist but they can only be extracted when fate is incorporated into the analysis.

Given that the clonal expansion phase (Figure 2a) induced partial differentiation (Figure 3a), we ran factor analysis again but only on single cells that remained in the most undifferentiated HSC cluster after this period. Among the 22 distinct CFMs that explained the dataset (Figure 4d-e), they corresponded to specific differentiation programs: a multilineage module (CFM3), an erythroid/mast cell module (CFM6), a myeloid module (CFM9), a mast cell module (CFM12), a lymphoid module (CFM14), and a cDC1 module (CFM20), among others (Supp Figure 6).

Several CFMs were similar between the earlier analysis of all HSPCs and this HSC-focused analysis. One unique to an HSC focus was a multilineage factor (CFM3). Importantly, this included genes (e.g. *ANGPT1* and *MECOM*) and markers (e.g. CD49f, CD81, HLA-DR) known in HSC biology, in addition to several that have not yet been described. Other CFMs mostly contained genes that do not have a described role for their corresponding lineage trajectory. One recurring surface marker that was present in several CFMs (multipotency CFM3, myeloid CFM9, lymphoid CFM14, DC1 CFM20, among others). While CD200 is known to be expressed in human HSC^31^, it was not previously associated with any specific HSC function. Intriguingly, CD200 was missing only from CFM6, the erythroid module. This led to the hypothesis that CD200 might be associated with HSC that can produce multiple cell types, except for erythrocytes.

In summary, factor analysis successfully uncovered hidden gene and surface marker modules correlated with fate outcomes. Notably, they were distinct from those optimal for WNN cluster classification and thereby represented hidden predictors of clonal human haematopoietic fate that only emerged upon integration of fate information through factor analysis. As CD200 was a hidden predictor of multipotency, and this marker has not been explored as a prospective HSC marker previously, we proceeded to test this marker functionally.

### CD200 demarcates multilineage HSC

HSCs are defined by their ability to repopulate the entire blood system upon transplantation. This capacity implies their ability to self-renew and to have multilineage potential. While HaemaTONIC is not necessarily a proxy for engraftment in humans or humanised mice, it can be used to test lineage potential and contribution over 7-8 weeks. Based on the existence of different clonal fate modules amongst HSCs of which CD200 was associated with multipotency we hypothesised that its expression would be heterogenous and demarcate multipotent HSCs from other HSCs. Analysis of CD200 expression in phenotypically defined human HSCs (CD34^+^CD38^−^CD90^+^CD45RA^−^) revealed heterogeneous expression patterns in both cord blood and G-CSF mobilized peripheral blood samples (Figure 5a-b). Interestingly, the percentage of CD200^+^ HSCs increased during the expansion phase, while the CD200^−^ HSCs declined, suggesting that the latter may have limited expansion capacity (Figure 5b). Comparative analysis with established HSC markers^32-34^ (ESAM, CD49f, CD66e) demonstrated that CD200-expressing HSCs represent a distinct population from previously characterized HSC subsets (Figure 5c).

To investigate potential functional differences between CD200^+^ and CD200^−^ HSCs, we isolated these populations from cord blood samples of multiple donors and evaluated them using our HaemaTONIC culture system. CD200^+^ cells demonstrated enhanced persistence in culture compared to CD200^−^ HSCs, suggesting superior self-renewal capacity. This observation was further supported by elevated frequencies of CD34^+^ progenitors in CD200^+^ cultures relative to CD200^−^ cultures (Figure 5d). Moreover, while CD200^+^ HSC numbers progressively decreased in culture, we were able to detect them up to 3 weeks when cultured in differentiation conditions (Figure 5d). Interestingly, CD200 expression persisted on sorted CD200^+^ HSCs, with near to 100% CD34^+^CD38^−^CD90^+^ HSC in the CD200^+^ condition retaining this marker in HaemaTONIC for the first 3 weeks of culture (Figure 5e).

While both CD200^+^ and CD200^−^ HSCs exhibited multilineage differentiation potential, we observed distinct patterns of lineage output. CD200^+^ HSCs generally produced higher cell numbers across all lineages. The most striking disparity was observed in DC development, where CD200^+^ HSCs generated substantial numbers of cDC1s, cDC2 and pDCs, while CD200^−^ HSCs failed to produce any (Figure 5f-g). Notably, certain populations (such as monocytes and neutrophils) were generated by both CD200^+^ and CD200^−^ HSCs but with distinct temporal patterns: CD200^−^ cells contributed to an earlier wave of production, while CD200^+^ cells dominated later-stage production. Interestingly, CD200^−^ cells demonstrated superior erythroid differentiation capacity, especially at later time points.

These findings conclude that CD200^+^ and CD200^−^ phenotypic HSCs represent two functionally distinct populations and supports the correlation of CD200 with multiple CFM in factor analysis. Moreover, CD200^+^ HSCs exhibit enhanced self-renewal capacity and sustained multilineage output in culture, whereas CD200^−^ HSCs show preferential erythroid differentiation potential, reminiscent of previously described HSC heterogeneity at a clonal level^15^.

### CD200 enrichment enhances DC1 manufacturing

Type 1 conventional DCs (cDC1s) have increasingly become a focus in immuno-oncology due to their crucial role in anti-tumor immunity^35^. These specialized antigen-presenting cells excel at cross-presentation and possess a unique capacity to activate CD8^+^ T cells, making them particularly adept in cancer immunosurveillance and a valuable target for therapy^36,37^. However, compared to the numerous GM-CSF driven monocyte-derived DC clinical trials, no dedicated cDC1 clinical trials have been completed. This is because, despite their therapeutic potential, strategies to efficiently generate human cDC1s in large numbers in stromal-free GMP settings remain limited. To our knowledge only one such published protocol exists, that involves a stem cell expansion phase, followed by a differentiation phase^38^. However, this method generated only 5% cDC1. Any method to increase purity of a DC1 product would have clinical utility.

We therefore investigated whether selecting for CD200-expressing HSPCs could enhance cDC1 production efficiency considering their preference for cDC1 generation. We evaluated this by enriching for CD200^+^ progenitors in the established protocol. CD34^+^ cells from three G-CSF mobilized donors were expanded, then MACS sorted into CD200^+^CD34^+^ and CD200^−^ CD34^+^ populations, followed by culture GMP protocol for 7 days^38^ (Figure 5h). Strikingly, CD200^−^CD34^+^ completely failed to generate any cDC1s, while CD200^+^CD34^+^ cells successfully produced them at high efficiency (Figure 5i). These findings suggests that enriching for CD200^+^ progenitors prior to DC1 manufacturing could significantly enhance the efficiency and purity of cDC1 by eliminating progenitors committed to alternative lineages.

Our results establish CD200 as a promising marker for multipotent HSCs and suggest its potential utility in optimizing cDC1 manufacturing for clinical applications. This discovery may have significant implications for improving the efficiency of cDC1-based immunotherapy approaches.

## Conclusions

Here, we utilise HaemaTONIC – our human multilineage haematopoiesis culture system – to dissect the genes that control fate decision in early progenitors. Previous work has already shown that human progenitors are heterogenous for both fate and transcriptome at the single cell level^7,12,15,25^. However, these studies did not directly link transcriptional programs with different clonal fates. Some attempts have been made in mouse haematopoiesis, where it was found that although lineage bias was determined early in the process, it was not entirely possible to predict fate based solely on transcription^18,22,39,40^. This is presumably because of some hidden features that are not detected through standard single cell transcriptomic analysis alone. Such features may be better uncovered when assessing epigenetics, metabolome or overall physiological features that can be inherited upon cell division to daughter cells in a way that dictates their fate.

scATAC-seq profiling of human HSPCs found some degree of lineage determination as early as HSC based on the differential chromatin accessibility of lineage bias transcription factors^41^. This is promising because it suggests that, unlike in mouse, in human haematopoiesis fate might be determined by molecular regulators that can be detected by scATAC-seq and scRNA-seq. Additionally, functional lineage tracing studies reveal heightened bias in human progenitors, particularly in adult haematopoiesis^11,12,15,42^.

Indeed, in our study, we found that factor analysis could predict fate from progenitor transcriptomic signatures. However, this signature was not composed solely by known lineage specific transcription factors or genes that define progenitor populations, but also by genes that could not have been discovered without linked clonal fate information. This shows that there are underlying molecular features that define fate that can be hidden by the stronger differentially expressed genes between different HSPCs. However, this work does not nullify the lineage trajectories that have been previously identified through scRNA-seq or ATAC-seq alone, but also adds novel genes that could play a role in fate determination that could not have been found otherwise.

This dataset reveals novel molecular signatures associated with specific lineage commitment and multipotency. As an example of the power of this approach, we identified CD200 as a key marker demarcating HSCs with enhanced multilineage potential and superior self-renewal capacity. Notably, CD200^+^ HSCs demonstrated significantly higher efficiency in dendritic cell production, particularly cDC1s, compared to their CD200^−^ counterparts. Given the growing importance of cDC1s in cancer immunotherapy applications, this finding has immediate translational relevance. Indeed, our discovery offers a practical solution for enriching CD34^+^ progenitor populations capable of generating cDC1s in clinical manufacturing protocols. We demonstrated that CD200^+^CD34^+^ cells were exclusively responsible for cDC1 production in an approved clinical protocol. Since isolation of mature cDC1s from culture systems presents significant challenges in clinical settings, our findings suggest an alternative approach: enriching for CD200^+^ progenitors at the start of culture to enhance final cDC1 purity. This strategy could improve the purity of cDC1 production for therapeutic applications.

Future work will focus on validating additional genes and surface markers identified through factor analysis as correlates of specific cell fate decisions. Our ultimate objective is to establish a comprehensive map of genes that regulate not only lineage commitment but also the temporal dynamics of cell production. This knowledge has significant translational potential, as it could be leveraged to optimize the production of specific cell types for clinical applications.

Limitations: There are some important limitations of this work. The culture system is not physiological, and involves exposing cells to a subset of cytokines, whose concentrations may not reflect those found in vivo, nor with the right timing of exposure to HSPCs during the differentiation process. Further, the absence of the natural niche environment may overlook effects that the niche could have on progenitor behaviour in vivo. Therefore, our culture system is a good proxy for what a cell can make and not necessarily for what it would make in an in vivo context. Another consideration is that our use of cord blood HSPCs may better represent foetal haematopoiesis rather than in adult. Any molecular regulators identified will have to be also tested on adult BM progenitors, considering the differences in potency between foetal and adult HSPCs^15^, or one would have to repeat these experiments using adult HSPCs.

## Methods

### CD34^+^ progenitor isolation

Cord blood samples were obtained from Murdoch Children Research Institute (MCRI). CD34+ progenitor isolation was performed following EasySep™ Human Cord Blood CD34 Positive Selection Kit II protocol (Stem Cell Technologies). In brief, PBMC were isolated by layering diluted blood on Ficoll-Paque™ PLUS density media and spinning down at 1200xg for 20 minutes. PBMC were then incubated with anti-CD34 Ab and a positive enrichment was performed using magnetics beads. Positive fraction was either put in culture or frozen in Bambanker medium (Fujifilm Biosciences) for future use.

### Cell culture

CD34^+^ progenitors were expanded for 7-10 days in 12 well plates in 1mL of StemMACS™ HSC Expansion Medium (Miltenyi Biotec) supplemented with StemMACS™ HSC Expansion Cocktail (Miltenyi Biotec) and 50 U/mL Penicillin-Streptomycin (Thermofisher). After expansion cells were either frozen or put in HaemaTONIC for differentiation (manuscript in preparation). For cDC1 culture using GMP media, CD34+ HPCs isolated from G-CSF mobilized blood cells were cultured with 1µM Stem Reginin1 (Stemcell technologies, Catalog # 72352), 50 µg/mL Ascorbic acid (Sigma), 2% human serum (Sigma), 100 ng/mL of rhFLT3L, rhSCF and rhTPO in Cellgenix® GMP DC medium for 7 days. To isolated CD200+ HPCs, expanded CD34+ cells were labelled with an APC-Cy7-conjugated anti-CD200 antibody (BioLegend). Subsequent enrichment for CD200+ cells was achieved through the application of Anti-Cy7 microbeads (Miltenyi Biotec) utilizing Magnetic-Activated Cell Sorting (MACS) technology. For comparative analysis, four distinct cell populations were obtained: enriched CD200+CD34+ HPCs, a control group of unfractionated CD34+ HPCs (No MACS), CD34+ HPCs processed through MACS in the absence of APC-Cy7 anti-CD200 antibody labelling (No CD200 Ab labelled), and the CD200-negative flow-through HPCs resulting from the MACS procedure (negative flow through). All isolated cell populations were washed with phosphate-buffered saline (PBS) and cultured in Cellgenix® GMP DC medium supplemented with 1µM Stem Reginin1 (Stemcell technologies, Catalog # 72352), 50µg/mL Ascorbic acid (Sigma), 2% human serum (Sigma) 800 U/mL rhGM-CSF, 1000 U/mL rhIFN-α and 100 ng/mL rhFLT3L for 7 days. Differentiated cell populations were characterized by Flow cytometry.

### Cell staining

Half the content of HaemaTONIC culture was harvested at the timepoint specified. For sorting, cells were stained first with a cocktail of Ab containing CD45-BV786, CD16-APC Cy7, CD14-PE Cy7, Clec9A-PE, CD1c-APC, CD123-biotin. Cells were then washed and samples were incubated with a cocktail of magnetic beads anti-PE, anti-APC, anti-Cy7 and anti-biotin (Miltenyi Biotec) for 10 minutes. Samples were separated into a positive (DC-Myeloid) and negative fraction using LS column (Myltenyi Biotec). The negative fraction was further stained with CD19-BV650, GlyA-BV510, CD7-BB700, cKit-APC.

### FACS analysis and sorting

Cells were stained in FACS buffer and analysed using either Symphony Analyser (BD Biosciences) or Aurora Analyser (Cyteck) using BD Diva or SperctoFlow software. Analysis was performed using FlowJo (v10). Cells were sorted on a BD FACSAria III cell sorter (BD Biosciences)

### SPLINTR virus production

SPLINTR plasmids were obtained from Dawson’s lab at Peter MacCallum Cancer Centre. HEK293T were plated in 10 cm petra dish at 3×10^6^ cells per dish in DMSO media (Gibco) with 10% FBS on day 1. On day 2 1.5ug of SPLINTR plasmid was mixed with 1.0ug of PsPax packaging plasmid and 0.5ug of VSVG envelop plasmid. The plasmid mix was incubated with in 200ul of OptimMEM medium with 15 ul of Fugene HD transfection reagent (Promega) for 15 minutes and then the plasmids were transfected on the HEK293T. From here after OptiMEM serum free medium (Gibco) was used. On day 3 a medium changed was performed and 2 media harvests were performed on day 4 and 5. Virus was concentrated using LentiX concentrator (Takara Bio) 100X and aliquots were frozen at −80 in OptiMEM serum free medium.

#### Human cell virus transduction

Cells were plated in 96 well plate U bottom at a maximum of 50K cells per well in 100ul of StemMACS™ HSC Expansion Medium (Miltenyi Biotec) supplemented with StemMACS™ HSC Expansion Cocktail (Miltenyi Biotec) and 50 U/mL Penicillin-Streptomycin (Thermofisher). SPLINTR lentivirus was added on the cell according to titration to get 20% transduction efficiency. Cell were spun down at 600 xg for 1 hr at RT and then incubate for 16 hr at 37 degree. Virus was washed the next day 3 times with FACS buffer 10% FBS.

### SPLINTR library processing and analysis

#### DNA extraction

DNA was extracted by incubating up to 10^6^ cells in 40 µL Viagen (Australian Biosearch, cat# AB302-C) with 0.4 mg/mL Proteinase K (Thermo Fisher Cat# 25530049). Cells were placed in a thermocycler for 2 hours at 55 °C, 1 hour at 85 °C and 5 minutes at 95 °C. After extraction DNA was stored at –20 °C and used for library preparation.

#### Barcode amplification and sequencing

110 µL of reaction mix was added to 40 µL of cell lysate in Viagen and then split into 2 reactions of 75 µL each. Reaction mix contained 1X Q5 High Fidelity polymerase (NEB, cat# M0492L), 0.1 µM of R1 primer mix (Table 2.3) and UltraPure™ DNase/RNase-Free Distilled Water (Thermo Fisher, cat# 10977015). PCR replicates were incubated in a thermocycler and a first PCR round was conducted (1 cycle at 98°C for 3 mins followed by 25 cycles at 98°C for 15 sec, 68°C for 15 sec, 72°C for 30 sec, then 1 cycle at 72 °C for 5 mins). 0.75 µL from the first PCR round was used as input for second round. In this round samples were indexed with 2 µL of xGen™ UDI 10nt Primer (IDT 10008053) in 20 µL reactions with 1X Q5 High Fidelity polymerase (NEB, cat# M0492L) (1 cycle at 98°C for 3 mins followed by 20 cycles at 98°C for 15 sec, 68°C for 15 sec, 72°C for 30 sec, then 1 cycle at 72 °C for 5 mins). Samples were pooled equivolume and the final pool was cleaned with 1.2X NucleoMag beads (Scientifix, cat# 744970-50) prior of sequencing. The samples were pooled equivolume and sequenced on the Illumina NextSeq 2000 P2 100 cycles kit with 87 cycles for Read1, 30 cycles for R2 and 10 cycles for index (1% Phi-X was added)

#### Barcode data analysis and visualization

SPLINTR barcodes were extracted from sequencing data using a C++ program written by Dr. Tom Weber. The barcode sequences were then filtered against a reference file and only barcodes present in the reference were kept for further analysis. Barcodes were then filtered for barcodes present in both PCR replicates. After filtering, the reads counts were normalised to 10^6^ for each sample (count per million, cpm). The cpm matrix was then used to make the plots using ggplt2 R package (Wickham, 2016).

### scRNA-seq and Total-seq

#### ScRNA-seq sample processing and sequencing

Cells were harvested form HameTONIC culture at day 7 and day 21 (Figure 1) or expanded barcoded progenitors were thawed in FACS buffer with 10% FBS (Figure 3). Cells were stained with Total-seq A Ab panel (BioLegend) diluted 1:5 in FACS buffer. After staining cells were washed for 3 times and were resuspended in FACS buffer containing Propidium Iodide (1:5000 dilution). Live cells were sorted on BD FACSAria III cell sorter (BD Biosciences). For Figure 3 SPLINTR barcode positive cells were sorted based on the expression of GFP, mCherry or BFP fluorescent proteins. 100K cells were loaded per capture on a Chromium Next GEM Single Cell 3’ HT Reagent Kit v3.1 within 1 hour from sorting (10X Genomics). After capture, antibody derived tag (ADT) and gene expression (GEX) libraries were prepared according to protocol 10X protocol. Transcriptome samples were sequenced on MGI G400RS PE100 cycles kits using 28 cycles for Read1, 100 cycles for Read2 and 10 cycles for indexes. ADT libraries were sequenced either on MGI G400RS PE100 cycles kits or on Illumina NextSeq 2000 P2 100 cycles kit using 28 cycles for Read1, 30 cycles for Read2 and 8-10 cyles for indexes. Both libraries were converted from Illumina to MGI libraries using MGIEasy Universal Library Conversion kit (MGI, cat # 1000004155).

#### scRNA-seq analysis

scRNA-seq data was aligned using 10x Genomics Cell Ranger 7.0.0 and analysed using Seurat R package^43^. After alignment, poor quality cells were filtered out from the dataset. We filtered out cells with more than 5% of reads that mapped to the mitochondrial genome (low quality or dying cells) or cells with less than 200 genes (empty droplets). The remining cells were demultiplexed using cellsnp-lite (https://github.com/single-cell-genetics/cellsnp-lite) and vireo (https://github.com/single-cell-genetics/vireo). After demultiplexing doubles and cells that were not assigned to a donor were also removed from the analysis. TotalSeq was also aligned using 10x Genomics Cell Ranger 7.0.0 and analysed using Seurat R package^43^.

### Factor analysis

For each barcode $b_i$, we aggregate the following quantities:

— $X^{(b)}_i$ a celltype by timepoint (8×5) matrix of its temporal output (in cell number equivalents)
— $X^{(e)}_i$ averaged gene expression among sequenced progenitors
— $X^{(m)}_i$ averaged marker expression among sequenced progenitors

We filter genes by variance and retain the top 10%, this leaves 3123 genes.

We retained 112 barcodes after filtering on HSC population.

$X^{(b)}$ is a $112 \times 8 \times 5$ matrix, while the gene expression $X^{(e)}$ and $X^{(m)}$ matrices are $112 \times 3123$ and $112 \times 163$. To learn the factors, we solve a fused non-negative (matrix, tensor) factorisation problem using a custom code (see details in Methods). Given a number of latent programs $k$, we learn a $112 \times k$ _activity_ matrix $U$ where $U_{ij} \ge 0$ quantifies the activity of program $j$ in the clone with barcode $i$. Each of the $k$ programs consists of coupled profiles of temporal clonal output, gene expression, and marker expression.

