## Supplemental Figures for "The hidden predictors of human haematopoietic clonal fate"

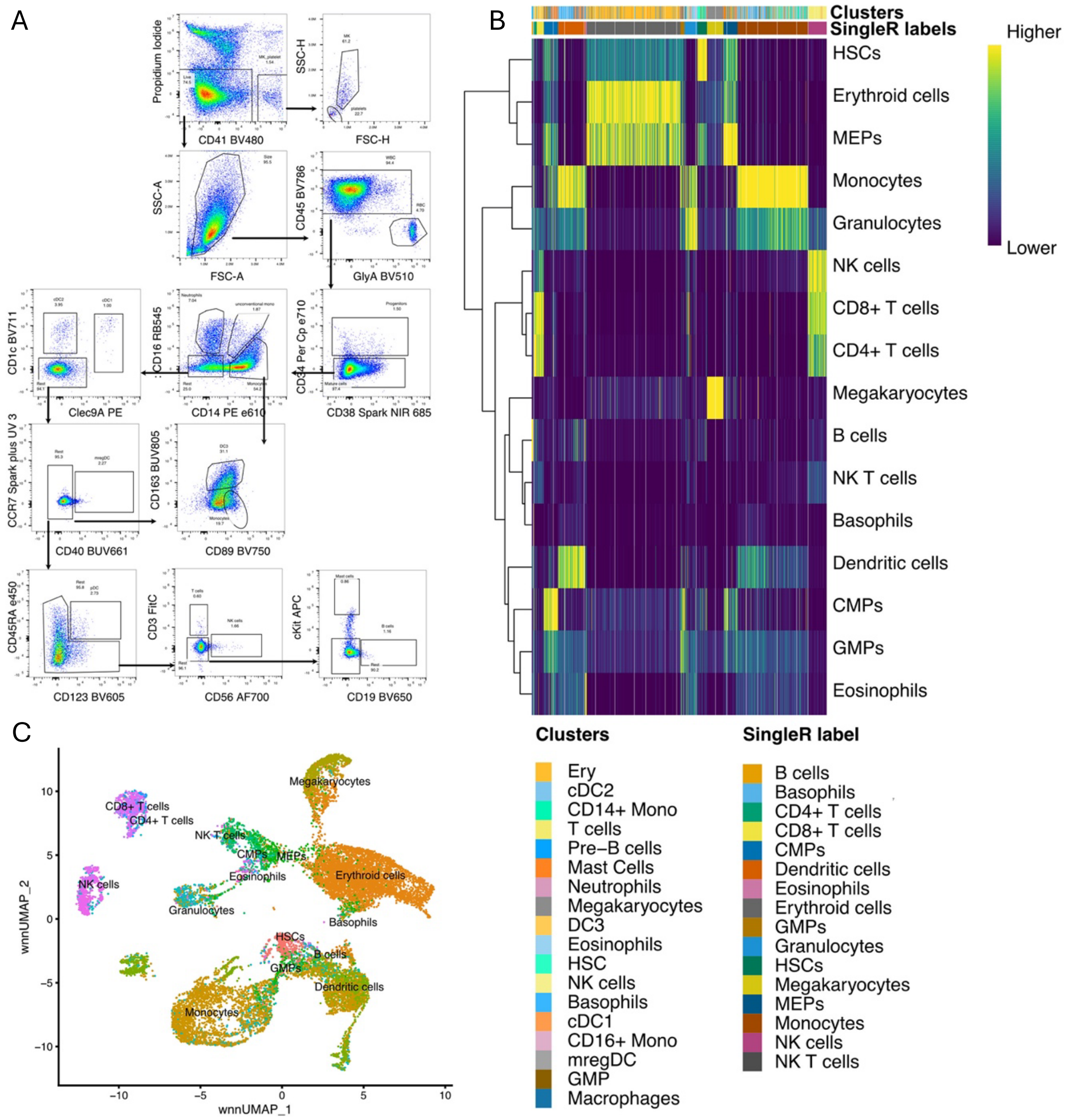


**Supp Figure 1: Gating strategy and cell type annotation with SingleR**

**a)** Gating strategy of HaemaTONIC culture using Spectral cytometry panel. **b)** Heatmap showing similarity score of mature populations generated in HaemaTONIC to human bone marrow dataset from and **c)** wnn umap of day 3 and day 21 HaemaTONIC timepoints annotated with SingleR

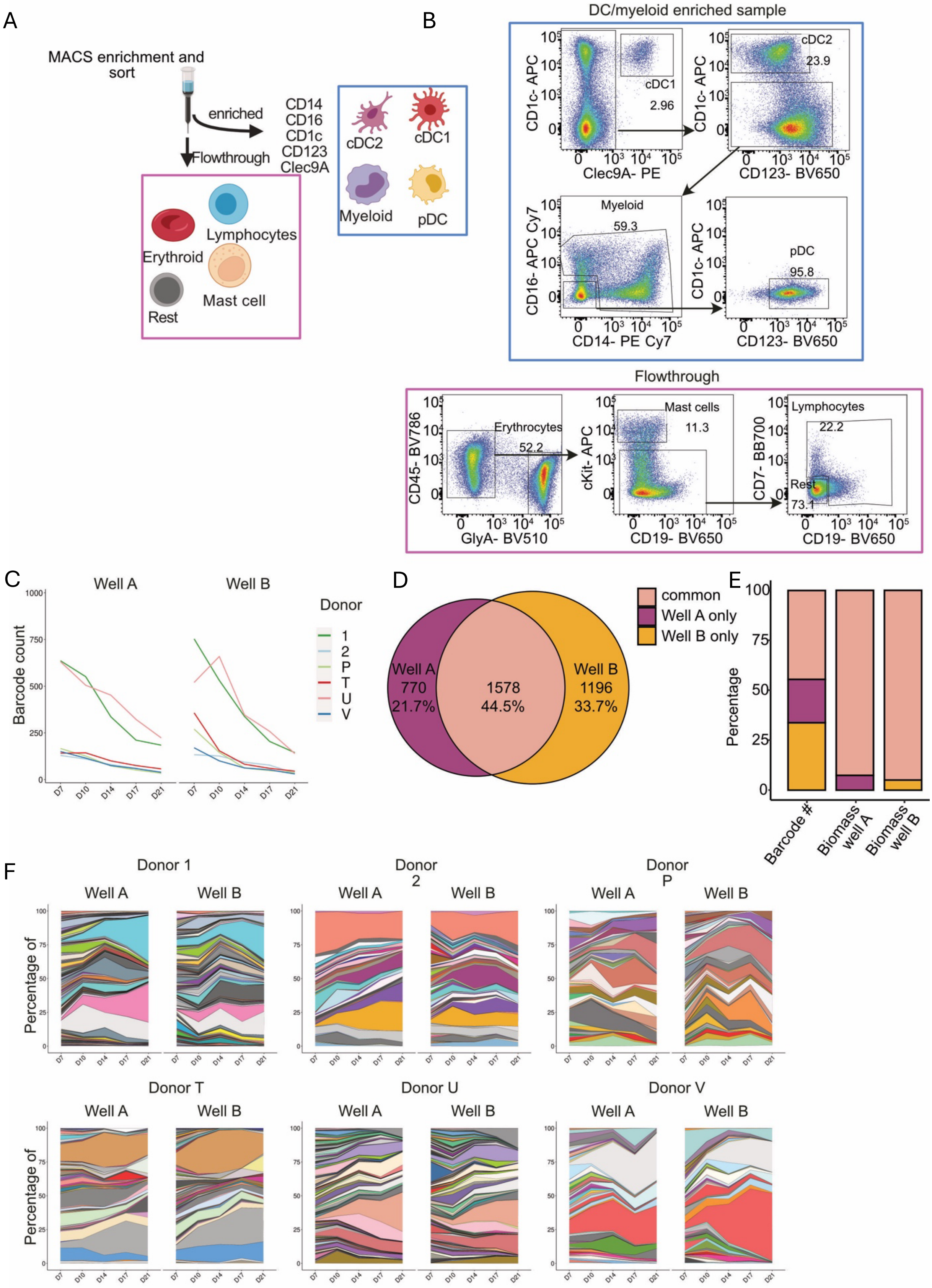


**Supp Figure 2: Clonal kinetics are conserved between wells**

**a)** MACS enrichment strategy to separate myeloid/DC fraction from rest and **b)** Gating strategy used to FACS sort the populations analysed in Figure 2. **c)** barcode number recovered from each donor for each timepoint analysed. **d)** Venn diagram showing number of barcodes present siter well A and B and number of barcodes in common between both. **e)** Histogram showing percentage of barcodes in common between wells and the percentage of the biomass that they produce in well A and well B. **f)** Area plot showing relative clone size of each barcode at each timepoint. The area under each barcode is proportional to the percentage of sequencing reads that the barcode got at each timepoint

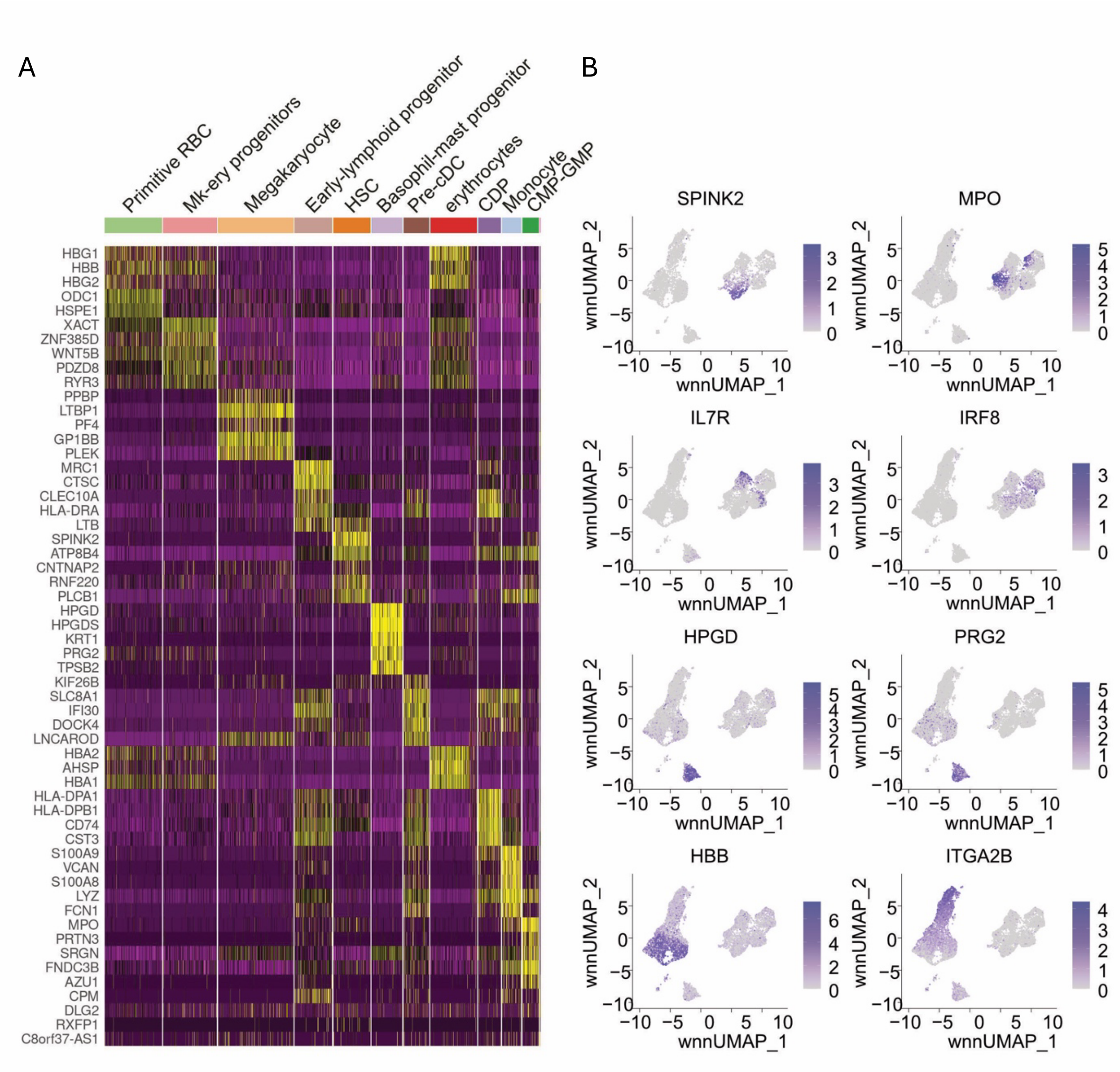


**Supp Figure 3: RNA clustering annotation**

**a)** Heatmap showing the top 5 differentially expressed genes for each cluster. **b)** Umap showing expression level of selected genes across all cells.

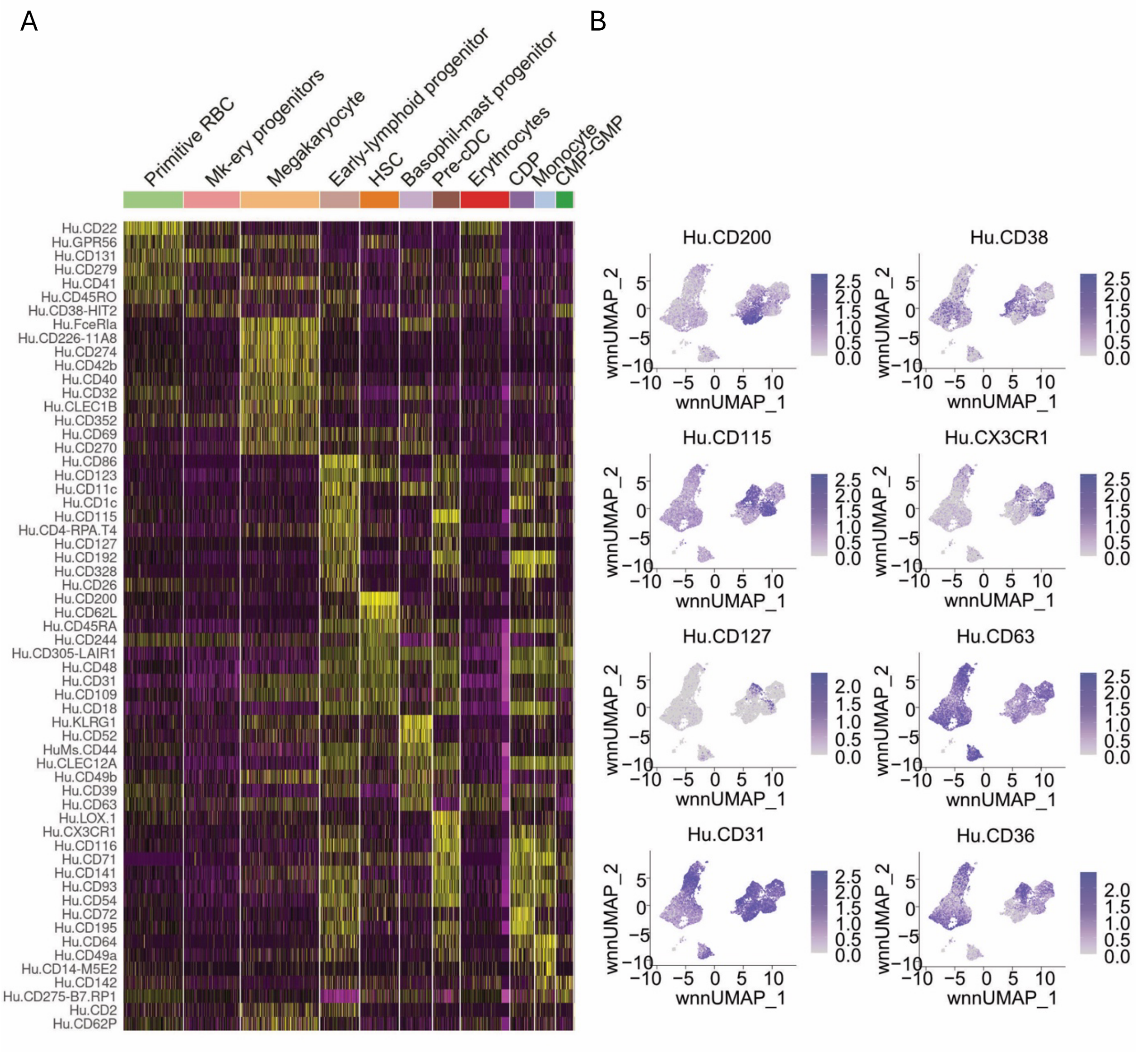

**Supp figure 4: Surface marker annotation**

**a)** Heatmap showing the top 5 differentially expressed Total-seq markers for each cluster. **b)** Umap showing expression level of selected Total-seq markers across all cells.

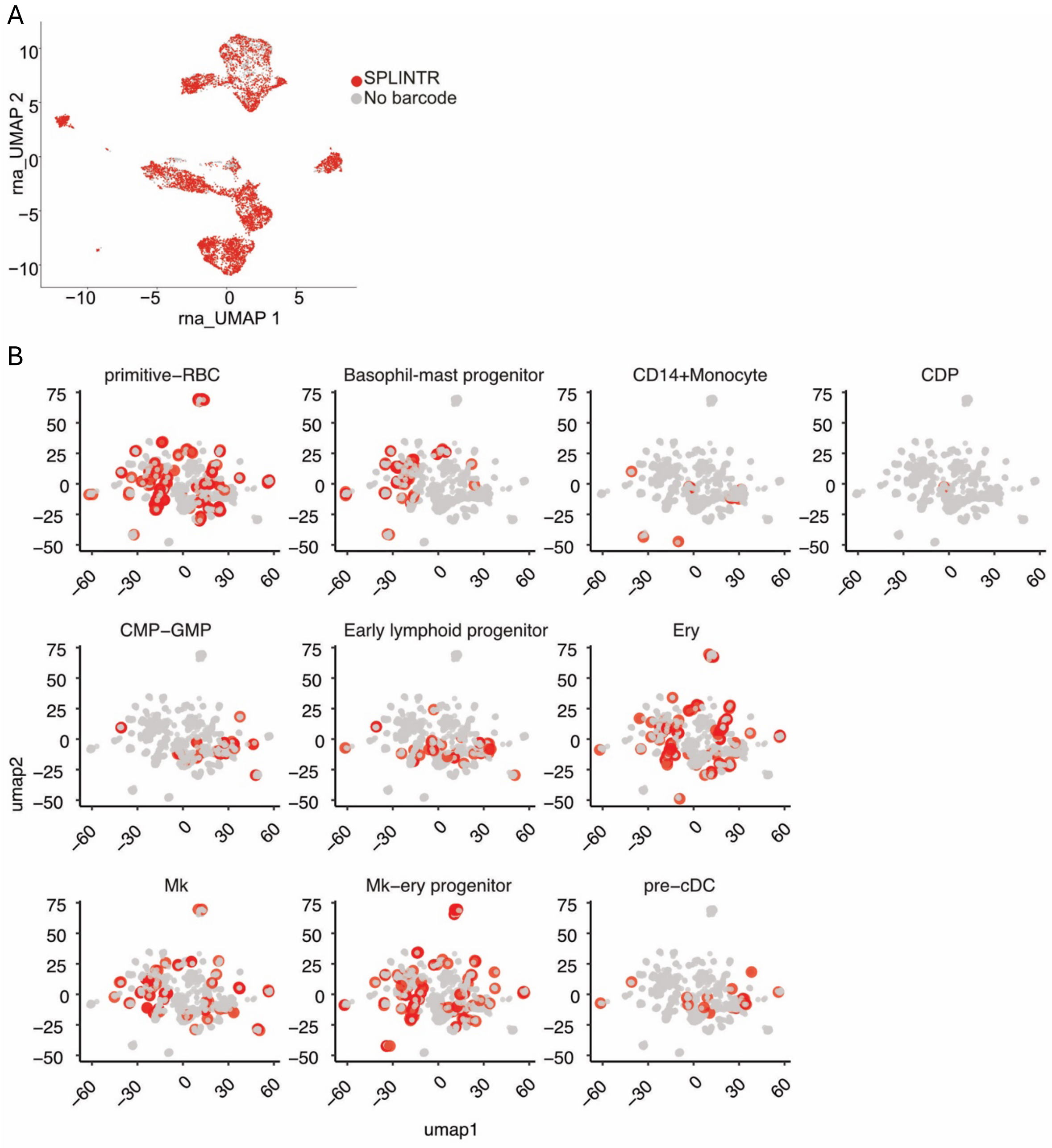


**Supp Figure 5: RNA based clustering does not correspond to fate clustering**

**a)** SPLINTR detection in scRNA-seq dataset. **b)** Overlay of cells from each Seurat cluster on fate UMAP according to their barcode

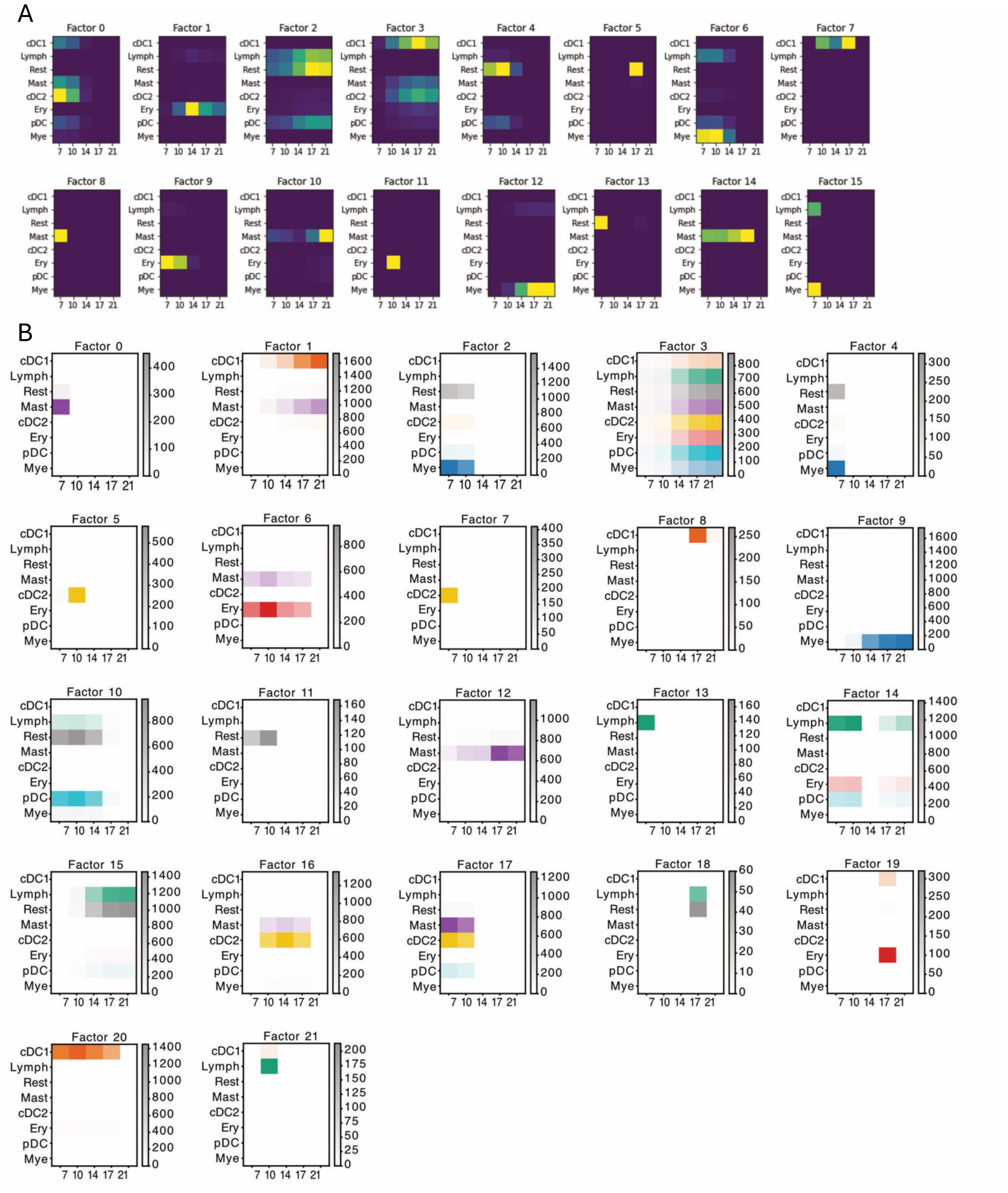


**Supp Figure 6: Factor analysis integrates fate and state**

**a)** Heatmaps showing the fate correlated with each factor for analysis performed with all progenitor cells types. **b)** Heatmaps showing the fate correlated with each factor for analysis performed with HSC cluster only
